# Point-localized and site-specific membrane potential optical recording by engineered single fluorescent nanodiscs

**DOI:** 10.1101/2021.04.21.440837

**Authors:** Asaf Grupi, Zehavit Shapira, Nurit Degani-Katzav, Shimon Yudovich, Shimon Weiss

## Abstract

Nanodisc technology was implemented as a platform for voltage nanosensors. A FRET-based voltage sensing scheme employing fluorescent nanodiscs and the hydrophobic ion dipicrylamine (DPA) was developed and utilized to optically record membrane potentials on the single nanodisc level. Ensemble- and single- nanosensor recordings were demonstrated for HEK293 cells and primary cortical neuron cells. Conjugation of nanodiscs to anti-GABA-A antibodies allowed for site specific membrane potential measurements from post synaptic sites.

## Introduction

Functional states of brain activity result from the correlated action of a large ensemble of neurons. With each neuron forming synaptic contacts with thousands of other neurons, the function of a neural circuit is an emergent property of these interactions, unpredictable by analysis of the functional properties of individual neurons[1]. While multi-electrode recordings have provided important insights from single (or small number of) neurons, they showed limited performance in analyzing dense local circuits or when signals from specific types of near-by neurons need to be distinguished. Moreover, membrane potentials are not uniform in time and space across the membrane of a single neuron and are likely to contribute to emergent properties of large neuronal systems[2]. An understanding of the functional properties of neural circuits requires dynamic recording of membrane potentials (and action potentials) from every neuron and its sub-cellular components.

For this reason, considerable efforts have been invested in developing optical recording methods that allow for simultaneous direct visualization of neuronal activity over a large number of neurons in a large field-of-view. Current state of the art voltage-sensitive dyes (VSDs)[3] and genetically encoded voltage indicators (GEVIs)[4, 5] are being used for neuroimaging at the ensemble level and in both in-vitro and in-vivo formats, however, their potential for neuroimaging is not fully exploited due to shortcomings such as alteration of membrane capacitance, phototoxicity and photobleaching[6]. More importantly, they do not allow for localized, site specific, membrane potential measurements.

Recently, voltage sensing nanoparticles (vsNPs) have been developed for non-invasive optical recording of membrane potential at the single particle and nanoscale level, at multiple sites, in a large field-of-view[7, 8]. In contrast to VSDs and GEVIs, vsNPs allow for nanoscale, single particle voltage detection. However, vsNPs still face challenges associated with functionalization, bioconjugation, stable membrane insertion, toxicity, and uniformity. For these reasons, the development of non-toxic and biodegradable voltage nanosensors would be highly advantageous.

Advances in the field of structural biology of membrane proteins have yielded nanodisc (ND) technology[9]. This technology is based on the self-assembly of engineered variants of apo-lipoprotein A and phospholipids into discoidal nanoparticles. Nanodiscs are relatively easy to produce, are composed of biological macromolecules and therefore biodegradable and non-toxic, can be modified and functionalized via their lipid headgroups or the protein side chains, and are capable of incorporating various types of lipids and lipophilic additives.

We identified NDs as a potential system for local membrane potential nanosensors by an established FRET based sensing mechanism[10]. We hypothesized that by incorporating several lipophilic (donor) dyes into a single ND and attaching it to a cell membrane, FRET based sensing of membrane potential could be achieved between the dyes in the ND and the non-fluorescent lipophilic ion dipicrylamine (DPA) which functions as a FRET acceptor. DPA partitioning between the inner and outer leaflet in response to membrane depolarization, leads to membrane potential dependent FRET changes

Exploratory testing of the utility of such NDs revealed complete quenching of donor dyes by the water-soluble fraction of DPA. This was resolved by incorporation of an additional fluorescent acceptor to the ND bilayer in order to provide a competing route for energy transfer, and using avidin as a steric barrier between the labeled NDs and the surrounding DPA molecules.

The performance of NDs as membrane potential sensors was tested and characterized by electrophysiology experiments conducted on HEK293 cells and primary cortical neurons, as well as site specific membrane potential measurements by antibody conjugated NDs. Outlook of the technology and future developments are briefly discussed.

## Results

In order to investigate whether donor labeled NDs can perform as membrane potential sensors via FRET to DPA, we labeled NDs with TopFluor® PE (TF), a BODIPY derivative labeled at the head-group of a PE lipid. This bright and stable lipid labeled fluorophore has a high extinction coefficient of 96,904 mol^−1^cm^−1^ in MeOH, a quantum yield (QY) of 0.9, and it incorporates spontaneously into the ND bilayer during the self-assembly process.

Initial experiments showed that the emission from labeled NDs adsorbed either to the plasma membrane of HEK293 cells or to a coverslip glass surface were completely quenched in the presence of 2 μM DPA in solution. This quenching is attributed to very high energy transfer efficiencies from TF to DPA molecules. The effect was reversible to some extent following several wash cycles and removal of DPA from solution thereby indicating that quenching occurs to either DPA molecules dissolved in solution, DPA molecules dissolved in the ND bilayer, or both.

In order to solve this detrimental effect, we employed two strategies aimed at (I) limiting the accessibility of water soluble DPA from the vicinity of solvent exposed dyes, and (II) reducing energy transfer efficiency to DPA (Fig. 1):

**Fig. 1.**
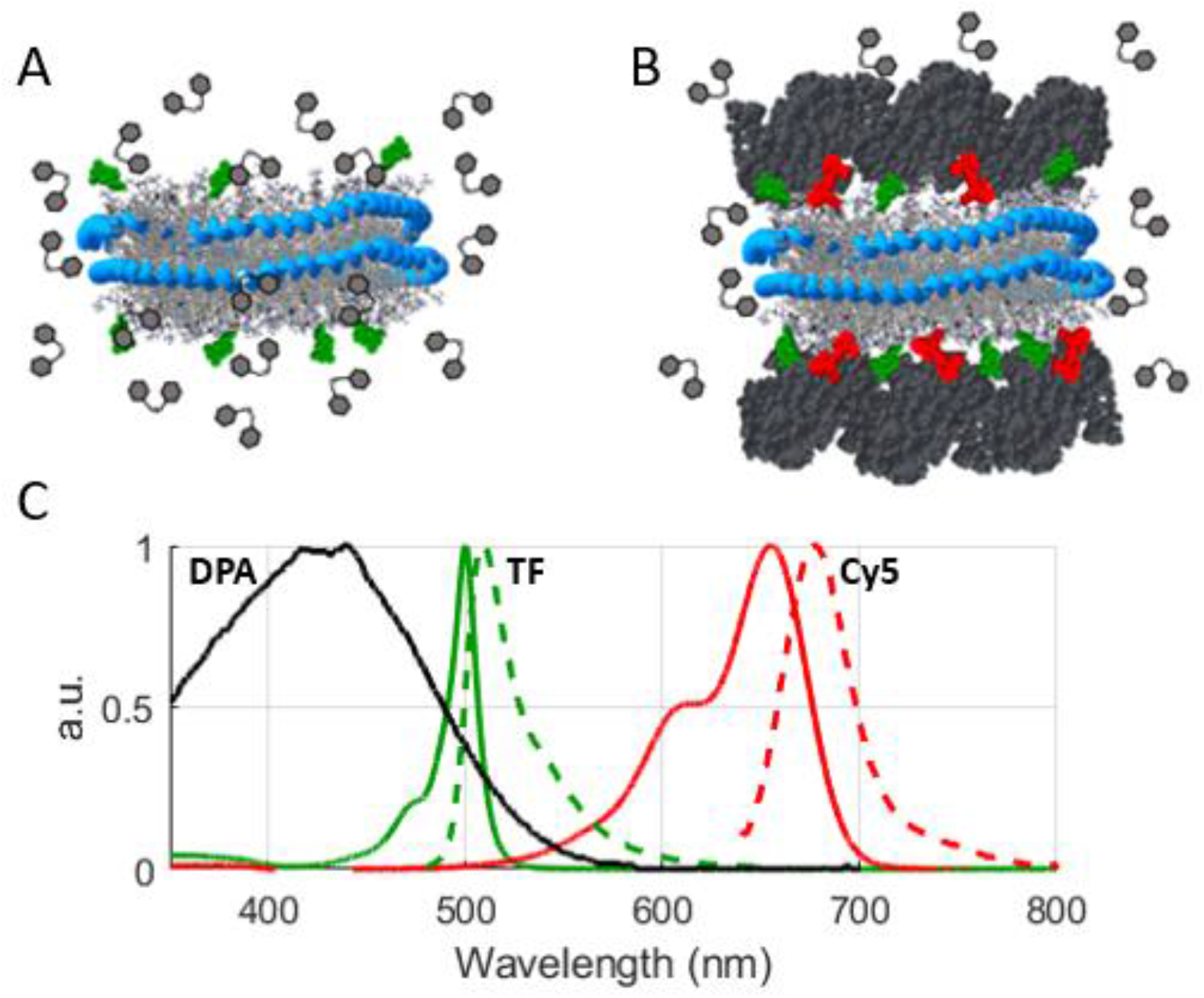
Scheme of functionalized NDs. **(A)** NDs labeled by TF (green) in the presence of DPA (gray hexagons). DPA is in both the solvent and the ND bilayer. **(B)** NDs labeled by TF and Cy5-PE (red). ND bilayer surface is capped by avidin (dark gray) which restricts the distance between DPA and TF. **(C)** The absorbance (solid) and emission (dashed) spectra of ND_TF_ (green) and ND_Cy5_ (red), and absorbance spectrum of 2 μM DPA (black). Absorbance and emission spectra were measured for ND_TF_ at ~10:1 TF:ND ratio and ND_Cy5_ at ~4.5:1 Cy5:ND. All spectra were recorded in PBS buffer at 20 °C. The overlap between the emission of ND_TF_ and the absorbance of ND_Cy5_ allows efficient energy transfer.

### (I) ND bilayer surface capping by biotin-avidin interactions

To limit the accessibility of DPA molecules to the vicinity of ND embedded dyes, biotinylated lipids were added to the ND composition. Avidin was conjugated to the solvent exposed biotin groups in the assembled ND, in order to form a steric barrier to DPA molecules. To ensure tight binding between avidin and the ND surface we used biotinylated lipids with a short spacer which permits only the solvent exposed biotin group to penetrate into avidin binding site, leaving no space for avidin translational motion. Furthermore, reaction parameters (biotinylated lipid concentrations, incubation time, avidin concentration, solution pH) were optimized to enhance the probability for multivalent binding of avidin to the ND surface[11, 12], thus increasing the contact area between avidin and the ND bilayer surface. This was achieved at a 40% mol ratio of biotinylated lipids to total ND lipids in the preparation mixture.

Avidin binding was confirmed by an observed hydrodynamic diameter increase from 13.1±2.5 nm to 27.2±4.6 nm, determined by DLS measurements. Considering that the hydrodynamic diameter of free avidin is 7.4±1.3 nm, the increase in diameter represents avidin binding to both facets. Incubation of biotin-free NDs with avidin under the same conditions resulted in large aggregates of >100nm in diameter, further supporting specific biotin-avidin association (Fig. S1). We note that nominal biotin concentrations lower than 40% were found to bind to avidin, manifested by a similar increase of hydrodynamic diameter of ND-avidin conjugates. However, these conjugates were not resistant to quenching by DPA, indicating non-multivalent avidin binding.

### (II) Incorporation of a fluorescent acceptor to NDs

In order to provide an efficient competing route to energy transfer from TF to DPA, we incorporated a second acceptor to the ND bilayer. We hypothesized that at high dye densities of TF and a second fluorescent acceptor, both confined to the ND bilayer, the average distance between the dyes would be short enough to produce high FRET efficiencies to effectively compete with excessive energy transfer to the soluble fraction of DPA molecules (which do not participate in membrane potential sensing). To that end, a Cy5 derivative conjugated to the headgroup of a PE lipid (Cy5-PE) was selected as a second acceptor to the TF donor. This dye incorporates into the ND bilayer during self-assembly, it has a high extinction coefficient of 220,000 mol^−1^cm^−1^ in DMSO and a QY of 0.51±0.05 when embedded into NDs. It is spectrally resolved from TF and has a Förster radius of 35.4 Å as the acceptor to TF.

### Optimization of labeled ND brightness

In order to determine the optimal number of TF dyes that would maximize the brightness of labeled NDs, the QY dependence on the number of TF and Cy5-PE was determined. This is necessary since the confined volume of the ND bilayer only allows for a finite number of dyes to be incorporated before reduction of brightness due to self-quenching and photon re-absorption begins to degrade the photophysical properties of the dyes. By preparing NDs with increasing number of labeled lipids, we observed that the brightness continued to increase upon the addition of labeled lipids until it reached ratios of 6:1 TF:ND for ND_TF_, and 4:1 Cy5-PE:ND for ND_Cy5_ (Fig. S2). Above 6 TF molecules per ND, and above 4 Cy5-PE molecules per ND, self-quenching greatly decreased the contribution of additional incorporated TF or Cy5 molecules to the construct’s brightness.

We note that the brightness of NDs labeled with TF at a 6:1 TF:ND ratio is very high compared to commercially available labels with a similar emission range used in biological applications. For example, the brightness of commercial streptavidin conjugated quantum dots (QD525) is only ~35% of ND_TF_ incorporating 6 TF dyes[13], and that of antibodies conjugated to Alexa 488 dyes at a working degree of labeling of 3 is only ~50% [14].

### Evaluation of labeled ND quenching by DPA

In the presence of 2 μM of DPA, which is a common working DPA concentration for FRET-based membrane potential measurements, the fluorescence from ND_TF_ with a ratio of 6:1 TF:ND was completely quenched. In order to test and compare the efficiency of the two approaches in reducing TF quenching by DPA, four labeled ND samples were prepared using 14 TF dyes per ND, which was enough to maintain sufficient fluorescence in the presence of DPA: (i) ND labeled with donor only at a molar ratio of 14:1 TF:ND (ND_TF_), and (ii) ND labeled with both donor and acceptor at a molar ratio of 14:6:1 TF:Cy5:ND (ND_TF,Cy5_). Both samples contained biotinylated lipids at a 40% molar ratio in the preparation mixture. Samples (i) and (ii) were diluted ×10 to a 5 mg/ml avidin solution in PBS pH 8.6 and incubated for 30 minutes at room temperature while stirring to produce samples (iii) ND_TF_-avidin, and (iv) ND_TF,Cy5_-avidin. All samples were filtered using a 0.22 μm syringe filter immediately before measurement. For photobleaching measurements of NDs adsorbed to a coverslip glass, the same samples were prepared with the addition of 3% DC-Cholesterol in order to add cationic charge to promote ND interaction with glass.

DPA quenching efficiency was evaluated by fluorescence lifetime measurements for samples in solution (Fig. 2), and by photobleaching kinetics imaging of labeled NDs adsorbed on a glass coverslip. From the photobleaching trajectories, we determined the average emission intensity for a single ND and the total emission time for each ND under constant illumination (survival time, see materials and methods). The quantification of these parameters allowed to evaluate the effect of DPA on the photophysical properties of different ND compositions, and is summarized in Table 1. The fluorescence lifetime for TF in ND_TF_, which was measured to be 2.23 ns in buffer without DPA, reduced to 0.8 ns with the addition of Cy5 in ND_TF,Cy5_ due to an expected high FRET efficiency between TF and Cy5-PE. However, the contribution of Cy5-PE to the photostability of TF in the presence of DPA is small, as shown in Table 1. This was also observed for glass adsorbed ND_TF,Cy5_, for which only ~8% of the NDs detected in buffer were detectable in the presence of 2 μM DPA, and with a ~50% and ~60% decrease in the mean intensity for TF and Cy5-PE, respectively.

**Table 1.** Photophysical properties of labeled NDs in buffer and in buffer with 2 μM DPA. ^1^Fluoresence emission was detected at a 520±15 nm and a 700±30 nm spectral range for TF and Cy5 emission, respectively. ^2^Fluorescence lifetime was collected for samples freely diffusing in solution using a 470 nm pulsed excitation. ^3^Survival time was defined as the characteristic total emission time of a single ND adsorbed to a glass surface. ^4^Mean ND intensity was defined as the average intensity for a population of NDs adsorbed to glass at the onset of illumination, with values normalized to the highest intensity measured. Adsorbed NDs were all excited at 488 nm using a constant illumination intensity. ^5^The proportion of detectable NDs per ROI in the presence of DPA relative to NDs detected in buffer. Evaluation was performed on the same glass surface.

|  | Sample ID | ND<sub>TF</sub> | ND<sub>TF,Cy5</sub> |  | ND<sub>TF</sub>-avidin | ND<sub>TF,Cy5</sub>-avidin |  |
| --- | --- | --- | --- | --- | --- | --- | --- |
|  | Emission Filter<sup>1</sup> | TF | TF | Cy5 | TF | TF | Cy5 |
| Buffer | Fluorescence lifetime (ns)<sup>2</sup> | 2.23 | 0.8 | 1.04 | 2.82 | 1.48 | 1.39 |
|  | Survival Time (s)<sup>3</sup> | - | 4.35 | 1.1 | - | 3.46 | 1.2 |
|  | Intensity (a.u.)<sup>4</sup> | ~0 | 0.31 | 0.99 | ~0 | 0.43 | 1.00 |
| Buffer + DPA | Fluorescence lifetime (ns)<sup>2</sup> | 0.36 | 0.47 | 0.38 | 0.96 | 0.65 | 0.89 |
|  | Survival Time (s)<sup>3</sup> | - | 6.77 | 4.4 | - | 3.9 | 3.2 |
|  | Intensity (a.u.)<sup>4</sup> | ~0 | 0.15 | 0.39 | ~0 | 0.12 | 0.28 |
|  | # NDs with DPA / # NDs without DPA<sup>5</sup> | ~0 | 0.05 | 0.08 | ~0 | 0.31 | 1 |

**Fig. 2.**
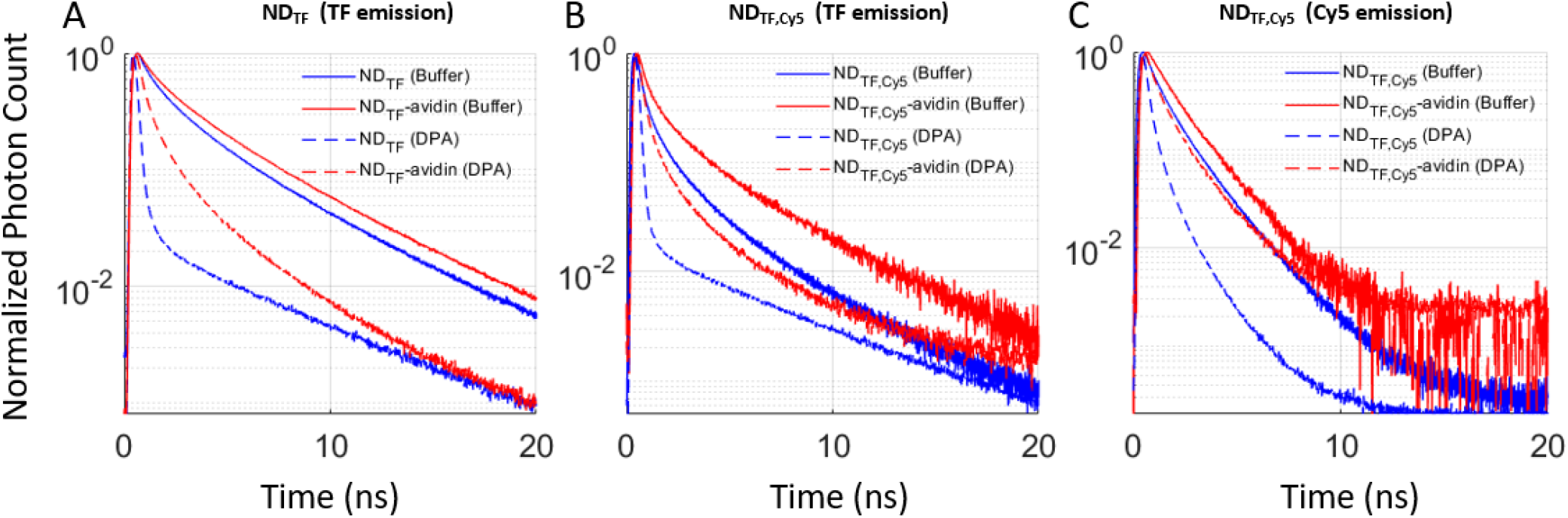
Fluorescence lifetime histograms of labeled NDs. Labeled ND samples were diluted into PBS to a final 30 nM in chambered coverslips and the fluorescence lifetime was recorded using a 485 nm laser excitation with a 520±10 nm emission filter for TF and a 700±30 nm for Cy5-PE. The normalized fluorescence lifetime decay curves are shown for NDs labeled with **(A)** TF only, **(B-C)** both TF and Cy5-PE, with **(B)** and **(C)** showing the fluorescence lifetime emission of TF and Cy5, respectively. Samples were measured in buffer (continuous line) and in buffer + 2 μM DPA (dashed line) for avidin-free (blue line) and avidin-conjugated (red line) constructs.

For the avidin conjugated samples, the fluorescence lifetime of TF for ND_TF_-avidin in buffer, increased from 2.23 ns to 2.82 ns, reflecting the change in the chromophore molecular environment. The lifetime for TF and Cy5-PE in ND_TF,Cy5_-avidin was longer compared to the avidin-free construct, but the photobleaching trajectories for both samples in buffer were similar. In the presence of DPA, lifetime decay curves of both TF and Cy5-PE in ND_TF,Cy5_-avidin, regain similarity to the decay curves of ND_TF,Cy5_ in buffer. This is also manifested in the properties of glass adsorbed ND_TF,Cy5_-avidin. The number of detectable TF and Cy5-PE in DPA compared to buffer, for glass adsorbed ND_TF,Cy5_-avidin, is 30% and 100%, respectively. Interestingly, the survival time of Cy5 was x3 fold higher compared to that measured in buffer. Moreover, while the recovery and survival time were higher, the mean intensity per ND was 20% for TF and 30% for Cy5 compared to the intensity recorded in buffer. The above leads to conclude a synergistic mechanism for sustainable dye emission in the presence of DPA, involving both surface capping by avidin and incorporation of an additional energy transfer channel for TF.

### Membrane potential sensing in HEK293 cells

The performance of ND_TF,Cy5_-avidin constructs as membrane potential sensors was first evaluated on the ensemble and single particle level using HEK293 cells as a model system. Cells were stained with 5 nM ND_TF,Cy5_-avidin, showing a high degree of staining (Fig 3B). The optical response of HEK293 labeled with ND_TF,Cy5_-avidin in a 2 μM DPA solution was measured by exciting TF at 470 nm, and detecting the emission of Cy5 at 665-725 nm, while voltage-clamping the cells to different values of membrane potential, stepping between −80 to 80 mV with 20 mV steps of 200 ms in duration, as shown in Fig. 3. The average optical response, defined as the relative change in fluorescent emission Δ*F*/*F*, from a total of five cells, was measured to be 33±6.4% per 120 mV (Fig. 3E).

**Fig. 3.**
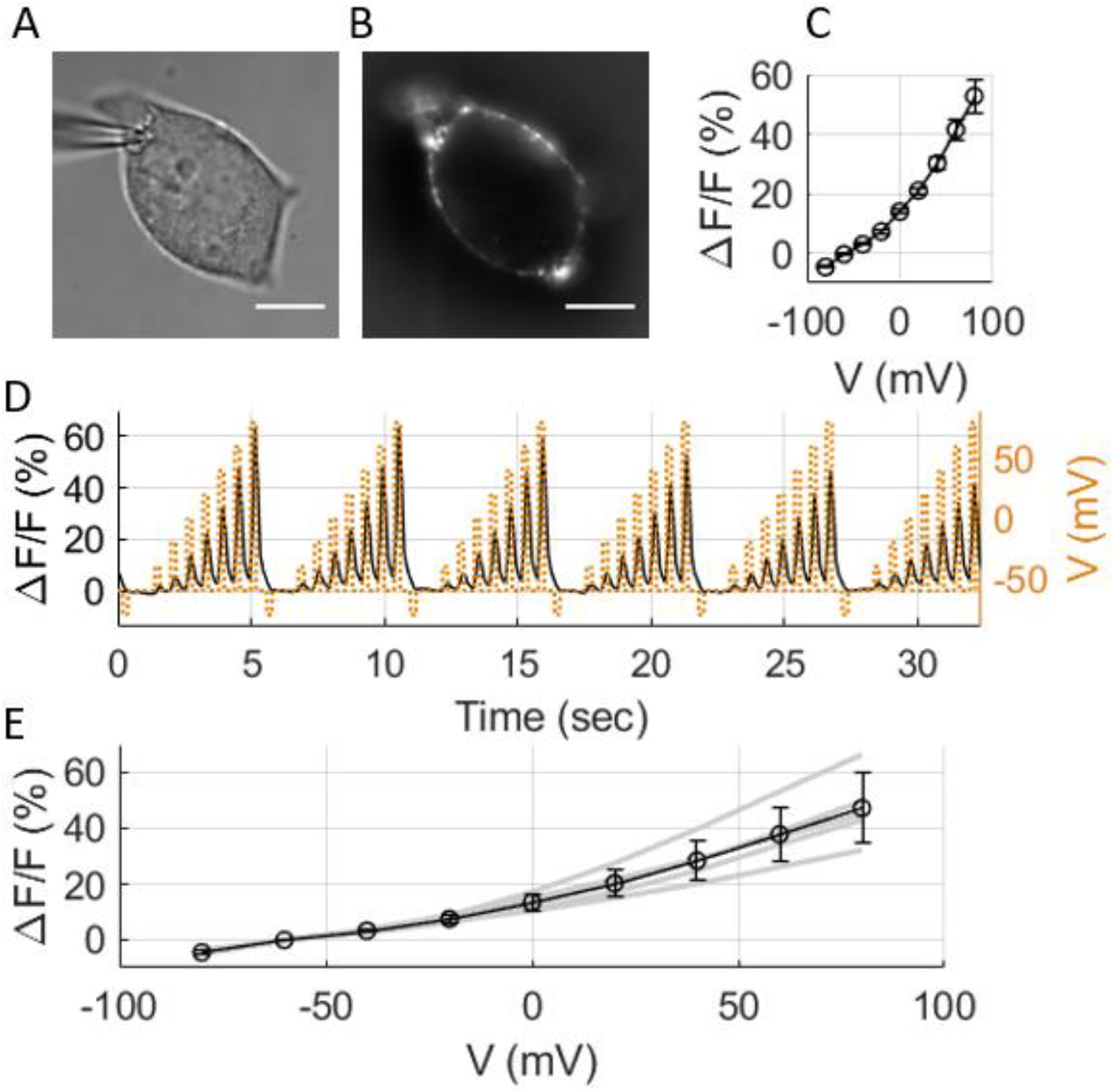
Fluorescence response of ND_TF,Cy5_ – DPA to changes of membrane potential in HEK293 cells. Cells were stained with ND_TF,Cy5_, and voltage-clamped in the presence of 2 μM DPA. **(A)** DIC image and **(B)** Cy5 fluorescence emission of a patch-clamped HEK293 cell. Scale-bar: 10 μm. **(C)** The averaged change in Cy5 fluorescence emission as a function of membrane potential, relative to the emission at −60 mV, averaging over 6 consecutive voltage steps. The voltage waveform consisted of varying voltage levels with 20 mV steps for 200 ms at each voltage level, and a 400 ms resting period of –60 mV. Optical recording was done at 10 Hz frame rate. **(D)** Error bars represent the standard deviation of the fluorescence change between consecutive voltage potential sweeps. **(E)** Fluorescence response curves of 5 individual cells (gray), calculated by averaging 6 consecutive voltage steps protocol, and the averaged response (black). Error bars represent the standard deviation of the average response values.

The response kinetics of ND_TF,Cy5_-avidin is not instantaneous with the change in membrane potential, and is limited to the detection of transient changes at the 10 ms scale and above, as further described in Fig. S4. We note that similar kinetics was observed when cells were directly stained with TF-PE labeled lipids. Since DiO-DPA FRET pair respond to changes in membrane potential at sub-ms timescales[10], it is likely that the fluorescent donor group of ND_TF,Cy5_ is also translocated by the change in membrane potential[15]. Another possibility for the slow response could be a slow mechanical translocation of the ND itself between membrane-tight and membrane-loose bound states (to be investigated in a future work).

### Membrane potential sensing in neurons

Primary cortical neurons were stained with ND_TF,Cy5_-avidin to achieve membrane labeling on the single particle level. Staining conditions were similar to those applied for ensemble labeling of HEK293 cells, however, due to lower membrane adsorption efficiency to neurons, single ND labeling was achieved. Neurons were voltage-clamped in a solution containing 2 μM DPA and the optical response of single ND_TF,Cy5_-avidin to changes in membrane potential was recorded. Membrane potential was alternated between −60, 0, and 60 mV. A representative example of the optical response of localized ND_TF,Cy5_-avidin staining on neuronal soma is shown in Fig. 4. The average response Δ*F*/*F* of 24 ND_TF,Cy5_ particles, measured from a total of five cells, to be 23.1±16% per 120 mV. We note that all the single fluorescent spots were responsive to changes in membrane potential, showing a change of fluorescent emission of at least 2% per 120 mV. These results clearly demonstrate the ability of ND_TF,Cy5_-avidin to probe neuronal membrane voltage changes from single localized spots.

**Fig. 4.**
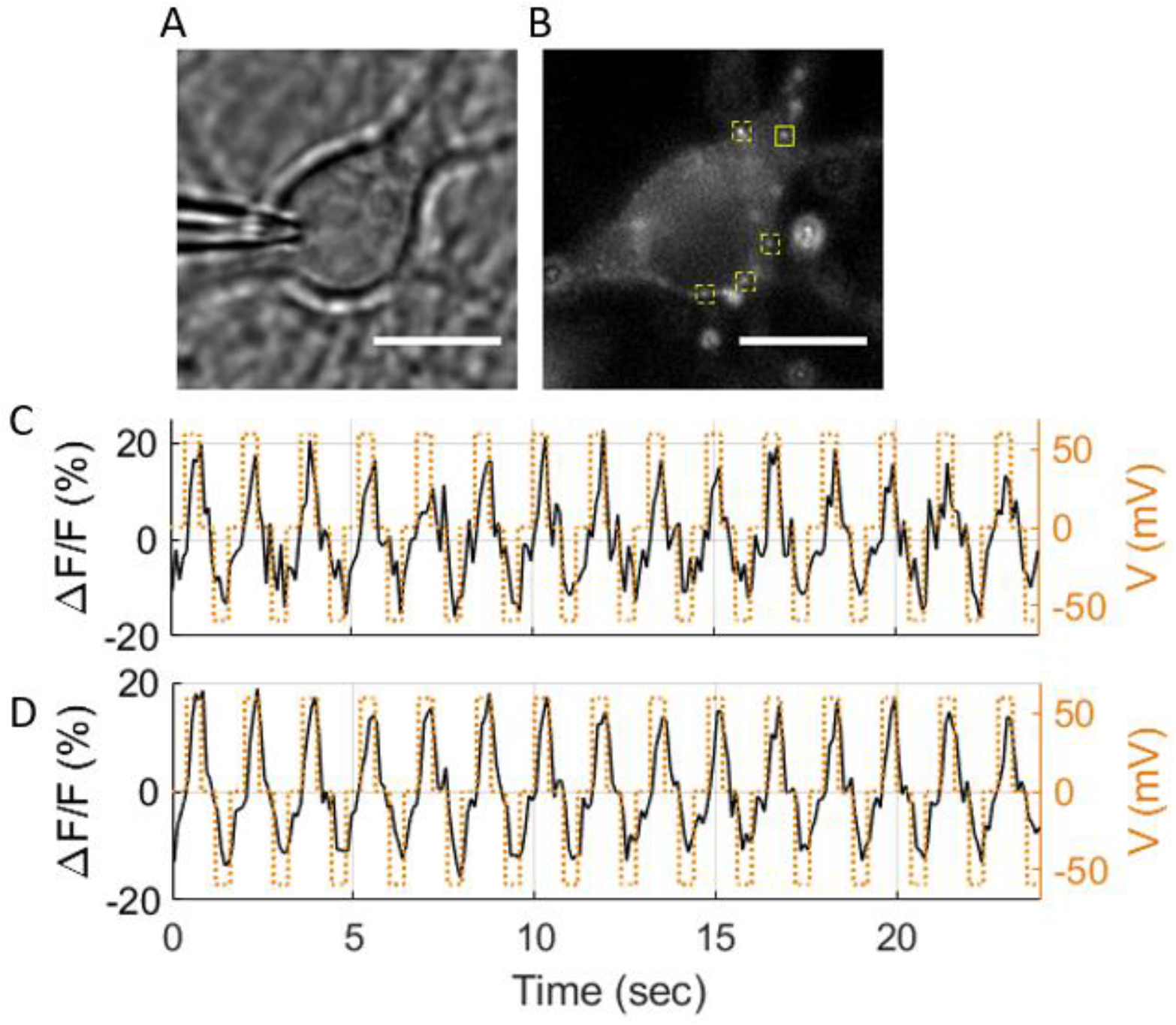
Fluorescence response of ND_TF,Cy5_ – DPA to changes of membrane potential in primary cortical neuron cells at low ND_TF,Cy5_ loading. Cells were stained with ND_TF,Cy5_, and voltage-clamped in the presence of 2 μM DPA. **(A)** DIC image and **(B)** Cy5 fluorescence emission of a patch-clamped primary cortical neuron. Scale-bar: 10 μm. The cells were clamped to a square voltage waveform consisting of 0, 60, 0, and −60 mV voltage levels, each 400 ms in duration. **(C)** shows the change in Cy5 fluorescence emission as a function of membrane potential from ND_TF,Cy5_ in the region inside the solid yellow square. Averaged response was 26.4 ± 5.2% per 120 mV. **(D)** The averaged fluorescence of five ND_TF,Cy5_ particles indicated by yellow squares, showing an averaged response of 27.1 ± 3.3% per 120 mV.

### Site directed membrane potential sensing in neurons

To demonstrate membrane potential sensing from specific neuron sites, we selected to target ND_TF,Cy5_ to GABA_A_ receptors (GABA_A_R). Depending on their subunit composition, GABA_A_R can be found on the post synaptic side of the synapse, and also on soma and dendrites [16, 17]. We used anti-GABA_A_ receptor γ2 (Ab-G) as the recognition element for conjugation to ND_TF,Cy5_. Ab-G targets the γ2 subunit, which is present in GABAergic synaptic junctions in the hippocampus, cerebellum, and most cortical synapses[18, 19]. To that end, ND_TF,Cy5_ were crosslinked to Ab-G via a hydrophilic linker to reduce potential undesired interactions between Ab-G and ND_TF,Cy5_ (ND-Ab) (figure 5A). Cell labeling was performed by incubation of ND-Ab with the neuron culture to allow for Ab-G driven membrane association, followed by a wash step and a short incubation with avidin to achieve ND bilayer capping by avidin and a strong membrane adsorption via the cationic charge of avidin.

**Fig. 5.**
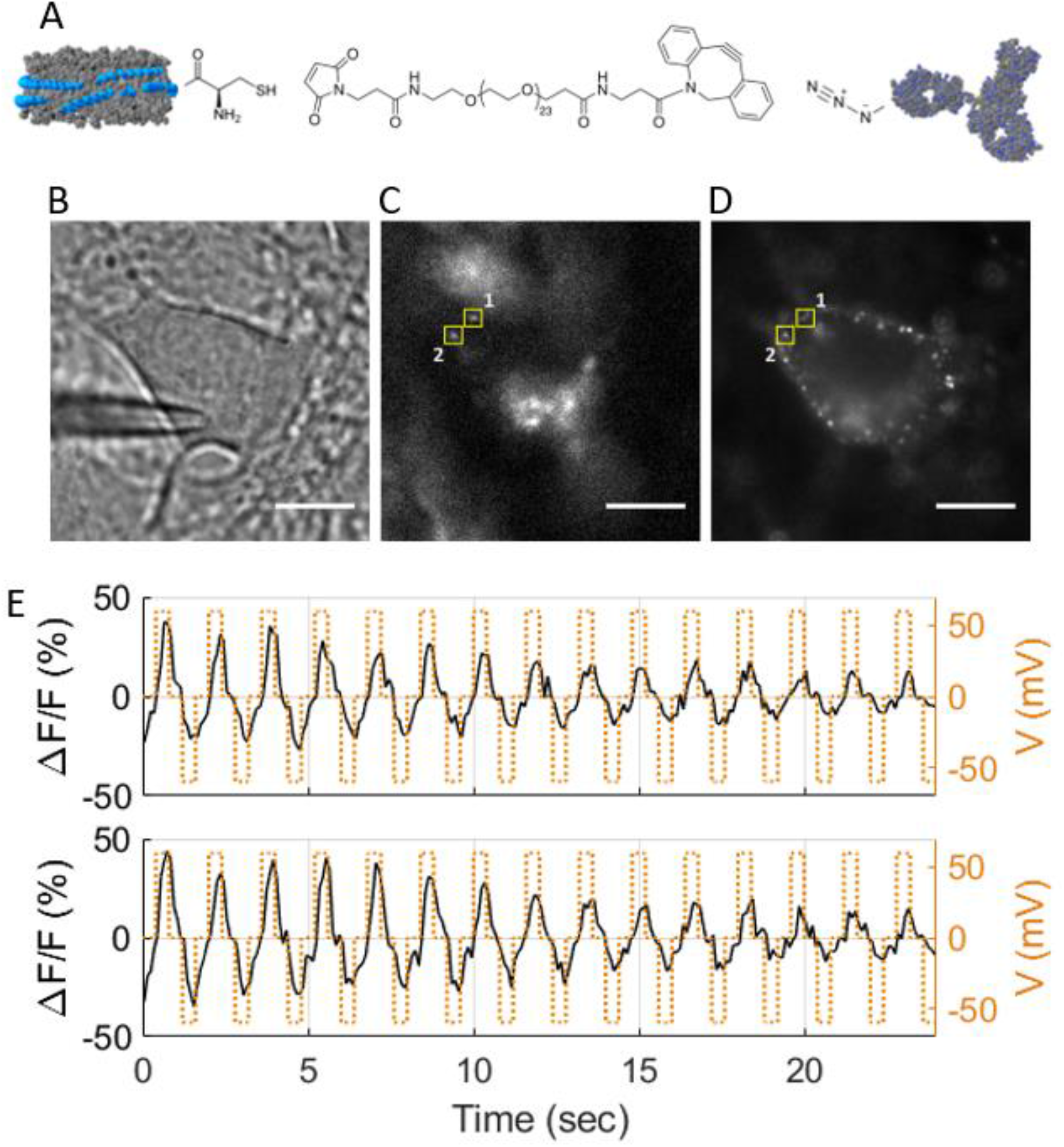
Single particle membrane potential optical recordings from ND_TF,Cy5_-Ab-avidin complex targeting GABA_A_R, and DPA. Cultured cortical neurons were transfected with mCherry-gephyrin and stained with ND_TF,Cy5_ conjugated to Ab-GABA and avidin. Cells were voltage-clamped in the presence of 2 μM DPA. **(A)** ND-Ab conjugation scheme, using Mal-PEG(24)-DBCO as a bifunctional linker. **(B)** DIC image of a patched cortical neuron, expressing mCherry-gephyrin and labeled with ND_TF,Cy5_-Ab-avidin. **(C)** and **(D)** show the emission of mCherry and Cy5, respectively. Scale-bar: 10 μm. **(E)** Shows the changes in Cy5 fluorescence emission as a function of membrane potential from the two ND_TF,Cy5_ (1 – top, 2 – bottom) in the regions inside the solid yellow squares. The average change in emission fluorescence was measured to be (1) 35.7%±14.4% and (2) 45.7± 20.1% per 120 mV.

To test for proper targeting, we transfected the cultured neurons with gephyrin fused to mCherry. Gephyrin is a cytoplasmic protein that accumulates at the post-synaptic complex of GABAergic and glycinergic synapses where it forms sub membranous lattices associated with postsynaptic clusters of GABA_A_R and glycine receptors, respectively[20]. Fig. 5 and Fig. 5S show colocalization and fluorescence responses to changes in membrane potential of transfected neurons stained with ND-Ab-avidin. Colocalizations between mCherry-gephyrin and ND-Ab-avidin (indicated by yellow squares), confirm successful targeting of GABA_A_R. We note that ND-Ab is expected to also bind to GABA_A_R γ2 which is colocalized with non-labeled endogenous gephyrin. However, negligible staining was observed when done with similar concentrations of ND without the conjugated antibody, supporting mainly specific binding of the conjugated ND.

We voltage-clamped neurons to a waveform containing three membrane potential levels, and recorded the fluorescence time trajectories from such single, targeted ND_TF,Cy5_, which appeared colocalized with mCherry-gephyrin.

Optical response to changes in membrane potential for two targeted ND_TF,Cy5_ shown in Fig. 5 were (1) 35.7±14.4% and (2) 45.7± 20.1% per 120 mV. As can be seen from Fig. 5E, the average error is mainly attributed to the photobleaching of the labeled NDs. In addition, the average response to changes of membrane potential was observed to be significantly heterogeneous, as seen from Fig. 5S. This heterogeneity could be a result of the heterogeneity of the NDs themselves, in terms of fluorophore labeling and avidin-binding, and to a lesser extent due to local differences in membrane potential occurring in whole-cell patch clamping of neurons.

## Discussion and Conclusions

We have demonstrated rational design and construction of multifunctional NDs for membrane potential sensing from specific sites in neurons. This was achieved by the unique chemical environment and precise self-assembly of NDs which affords modularity and custom modifications of ND composition. The capability to pack a large number of lipophilic dyes resulted in high brightness. We suggest that the performance of fluorescent NDs can be further improved by optimization of lipid composition and by inclusion of other photostabilizers such as triplet state scavengers to create self-healing nanoparticles, and that such improvements could extend the general applicability of labeled NDs to other biological applications.

Compared to voltage sensing QDs and NRs, labeled NDs provide an ideal solution for *in-vivo* applications since they are non-toxic, biodegradable and contain functional groups for bio-conjugation either as amino acid headgroups of the membrane scaffold protein or functionalized lipid head groups incorporated into the ND. Moreover, labeled NDs may provide an environment-friendly and competitive solution to toxic fluorescent nanoparticles currently employed in other biomedical imaging applications (beyond voltage sensing).

Here we applied a FRET based voltage sensing scheme based on a stationary, membrane adsorbed ND (donor) and DPA (acceptor), a hydrophobic ion that translocated across the membrane in response to membrane potential changes. The initial motivation to apply this sensing scheme was that donor dyes can be non-selectively incorporated into the ND bilayer, i.e. voltage sensing does not depend on a facet selective labeling of ND bilayer. A major obstacle for utilization of this sensing scheme was the high sensitivity of TF labeled NDs to quenching by DPA molecules in the buffer solution. We minimized this sensitivity by introduction of a competing acceptor to the ND bilayer which further improved by capping the ND surface with avidin and allowed to restore ND emission in the presence of DPA in solution. These strategies convey two additional advantages. The first, is that background emission is much lower for Cy5 emission compared to TF emission and the second is that the positively charged avidin also promotes adsorption to cell membranes.

We note that the addition of a competing de-excitation transition route to TF through FRET to Cy5 may decrease the overall sensitivity of the TF-DPA pair to changes in membrane potential. The average emission intensity of TF *F*_*TF*_ through direct TF or indirect Cy5 emission channels is proportional to *k*/*k*_*T*_, where k is the radiative transition rate of TF or the FRET rate to Cy5, and *k*_*T*_ is the sum of all radiative and non-radiative de-excitation transition rates of TF. Assuming the only voltage-dependent component of *k*_*T*_ is the non-radiative transfer of energy to DPA *k*_*DPA*_(*V*), the relative change in emission due to small changes in membrane potential is Δ*F*/*F* = −Δ*k*_*DPA*_(*V*)/*k*_*T*_. Therefore, an additional de-excitation route to TF inevitably reduces its sensitivity to changes in membrane potential. However, since the autofluorescent background of live cells is considerably weaker in Cy5’s red spectral range compared to TF’s green spectral range, imaging NDs that contain a TF and Cy5 FRET-pair results in a better optical signal to noise ratio for the detection of changes in membrane potential, especially at the single particle level.

Neuron and HEK293 cell membrane labeling with NDs was achieved by staining with NDs pre-formed with avidin bound to the ND bilayer. Since the positively charged avidin drives adsorption to the cell membrane, it is not likely that ND_TF,Cy5_-avidin binds parallel to the membrane since the distance gap by avidin should substantially reduce TF energy transfer efficiencies to DPA. It is therefore plausible that in this case, NDs bind perpendicular to the membrane.

A limiting factor for using the ND_TF,Cy5_ construct reported here for action potential sensing is its low temporal response. Using ND_TF,Cy5_, we were able to measure changes in membrane potential occurring at down to 10 ms, while action potentials are on the sub-ms time scale. One possible explanation is the observation that the temporal response measured for HEK293 labeled by TF is also slow and therefore it is only a matter of dye properties. To test this hypothesis, we performed voltage imaging of cells labeled by DiO and cells labeled by ND_DiO_ and found that while DiO showed a fast temporal response it was not the case for ND_DiO_, which points to ND related hindered temporal response. A similar slow temporal response was observed also with fluorescent polystyrene beads acting as membrane potential sensors on the nanoparticle level (DOI: XXXX). The observation of slow temporal response for two different types of nanoparticles points towards a similar underlying mechanism related to membrane adsorption. The reason for this slow temporal response of nanoparticles and for the slow response of the TF/DPA system is the subject of future studies.

Furthermore, incorporation of VSDs into NDs (that do not rely on FRET) would be the preferred system of choice as it combines the credibility and proven performance of VSDs together with the feasibility to perform voltage imaging. However, VSDs must be incorporated to one of the two membrane leaflets to be functional (otherwise they cancel the signals), which calls for facet selective labeling of NDs. Future developments include facet selective inclusion of VSDs into nanodiscs to provide a single component sensor.

## Supporting information

SI - Point-localized and site-specific membrane potential optical recording by engineered single fluorescent nanodiscs

## Contributions

AG conceived and planned this study, designed and prepared the materials, performed experiments and analyzed data. NDK, SY and ZS performed electrophysiology experiments and analyzed data. NDK and ZS planned, designed and performed the biological aspects. SW designed and planed this study. All co-authors participated in writing the manuscript.

## Acknowledgments

We thank Prof. Dan Oron and Dr. Miri Kazes help with quantum yield measurements and for helpful discussions and Eugene Brozgol and Prof. Yuval Garini for help with lifetime measurements. This work has received funding from the European Research Council (ERC) under the European Union’s Horizon 2020 research and innovation program under grant agreement No. 669941, by the Human Frontier Science Program (HFSP) research grant RGP0061/2015, by the BER program of the Department of Energy Office of Science grant DE-FC03-02ER63421, by the STROBE National Science Foundation Science & Technology Center, Grant No. DMR-1548924, by the Israel Science Foundation (ISF) Grant 813/19, and by the Bar-Ilan Research & Development Co, the Israel Innovation Authority, Grant No. 63392.

## Data and code depository

Raw data of all fluorescent imaging from all voltage measurements, TCSPC for this paper can be found here: ((https://zenodo.org, DOI: 10.5281/zenodo.4704309). Home-written code (MATLAB®) for extraction and analysis of recorded electrophysiological optical trajectories is available upon to request.

## Materials and methods

### Materials

Lipids (DMPC, TopFluor PE, Biotynil-PE (870285), Cy5-PE and DC-Cholesterol) were purchased from Avanti Polar Lipids. anti- GABA_A_ receptor γ2 IgG (cat#: 224004) was purchased from Synaptic Systems, Antibody azido modification click (SiteClick) from Thermo Fisher, and all other materials were purchased from Sigma Aldrich at the appropriate purity level.

### Plasmid extraction and purification

pMSP1E3D1 (Addgene plasmid #20066;http://n2t.net/addgene:20066; RRID:Addgene_20066) and pMSP1E3D1_D73C (Addgene plasmid #39328;http://n2t.net/addgene:39328;RRID:Addgene_39328) were a gift from Stephen Sligar. Plasmids in a bacterial stab were isolated by streaking bacteria onto an agar plate containing 50mg/ml kanamycin, followed by overnight incubation at 37 °C and selection of a single colony for overnight inoculation in 10ml of LB/kanamycin at 37 °C. Cells were harvested by centrifugation at 6000xg for 10min and the pellet was re-suspended in LB/kanamycin for plasmid extraction and purification using a GenElute plasmid miniprep kit (Sigma).

The plasmid was transformed into BL21(DE3) e. coli strain. Single colonies isolated from LB-agar/kanamycin plates were inoculated for 6 hours at 37 °C, harvested by centrifugation at 6000xg for 10 minutes and re-suspended with fresh LB/kanamycin. The suspension was diluted 1:1 with glycerol to a final 50%, mixed and dispensed to aliquots and stored in −80 °C.

### MSP expression

BL21(DE3) bacteria transformed with MSP1E3D1 or MSP1E3D1_D73C plasmids were slowly thawed on ice and 50 μl of the thawed bacteria was transferred to 10ml LB broth containing 50mg/ml kanamycin. The suspension was inoculated at 37 °C with shaking at 220 RPM until an OD_600_ of 0.8-1 was reached. The cell suspension was centrifuged at 6000xg for 10 minutes and the pellet was re-suspended in 10 ml of fresh LB/kanamycin solution. 5ml of the suspension were transferred to 500 ml of EZbroth/kanamycin in a 2L flask for shaking at 37 °C, 220 RPM until an OD of 2.5 was reached. Then, 500 μl of a freshly prepared 1M IPTG stock were added to the cell suspension for a final 1mM IPTG for initiation of protein expression. Cell culture was further incubated at 37 °C for 4 hours. Cells were harvested by centrifugation at 6000xg for 10 minutes. cell pellet was collected and stored in −80 °C.

### MSP purification

In a typical preparation, ~10 g of cell pellet were thawed on ice and re-suspended in 90 ml PBS 20mM. After complete re-suspension, 10 ml of 10% Triton X-100 were added to a final 1% concentration with stirring. Deoxyribonuclease was added to a final 2.5 μg/ml and the solution was left to stir for 20 minutes on ice. Cells were lysed by probe sonication using a 3mm probe (Sonics, VCX 130) (three one-minute rounds at 40% intensity or until the suspension changed color to dark brown). The suspension was centrifuged at 11,000xg for 45minutes in 50 ml centrifuge tubes to remove cell debris and finally filtered by a 0.22um syringe filter.

The solution was loaded on a Ni-NTA column pre-equilibrated with 40mM phosphate buffer. Column outlet was connected to a UV flow cell (Hellma, 0.1mm path length). UV absorbance of the elution volume was monitored at 280 nm by an Ocean Optics USB4000 spectrophotometer. The column was washed with 100ml of each of the following: (i) 40mM Tris/HCl 300 mM NaCl 1% Triton X-100 pH 8, (ii) 40mM Tris/HCl 300mM NaCl 50mM Na-Cholate 20mM Imidazole pH 8 (iii) 40mM Tris/HCl 300mM NaCl 50mM Imidazole pH 8. Finally, MSP was eluted by column wash with 40mM Tris/HCl 300mM NaCl 400mM Imidazole pH 8. MSP containing fractions were dialyzed against 20mM Tris/HCl 100mM NaCl pH 7.4 at 4 °C, following sample concentration using a 10 kD ultrafiltration device (Amicon, Millipore) and a buffer exchange to water. Samples stored at −80 °C at 200 μM.

In purification of the cysteine engineered variant MSP1E3D1_D73C, 5 mM b-mercaptoethanol was added to all the solvents used in the purification process.

#### ND preparation

ND composition was calculated to give a final ratio of 150:1 lipids:MSP with DMPC as the main lipid constituent. In a typical preparation, stock solutions of lipids stored in chloroform were mixed to yield the specific mol ratio composition. The solvent was evaporated using a vacuum concentrator (Labconco) for 20 minutes at 60 °C. The dried film was thoroughly dissolved in 8 μl of 20 mM Tris pH 7.4, 50mM sodium cholate and 100 mM NaCl and incubated for 15 minutes at 40 °C with agitation. To the lipid solution, 5 μl of a 200 μM MSP in water were added to give a final MSP concentration of 70 μM. The sample was incubated for 15 minutes at 40 °C with agitation. The solution was added to 7.5 mg of damp adsorbent beads (Amberlite XAD-2) pre-heated to 40 °C and incubated at 40° C with agitation for 1 hour. The sample was diluted with 100 μl of PBS, filtered by a 0.22 μm PES syringe filter and stored at 4 °C until further use. Sample purity was verified by size exclusion chromatography on a Superdex S-300 10/300 GL column (Pharmacia) and by DLS (for samples without Cy5-PE). The molecular weight of the purified MSP1E3D1 scaffold protein variant was determined by mass spectroscopy to be 32,601 D in agreement with the calculated molecular weight. The hydrodynamic diameter of a typical ND preparation was 12.9 nm as calculated by the retention time on the calibrated size exclusion column and 15±5 nm as determined by dynamic light scattering measurements on a Malvern Zetasizer Nano instrument. QY of fluorescent NDs was measured using a commercial quantum yield spectrometer (Quantaurus-QY C11347, Hamamtsu).

#### ND-Avidin complex preparation

Avidin was dissolved in PBS pH 8.6 to a final concentration of 5 mg/ml. 20 μl NDs incorporating biotinylated lipids were added slowly to 180 μl avidin solution under stirring and the solution was stirred for 20 minutes in the dark. The sample was filtered by a 0.22 μm syringe filter and centrifuged at 20K RCF, 4 °C for 15 minutes, and used immediately.

### ND conjugation to Ab

Site specific conjugation of NDs to anti- GABA_A_ receptor γ2 IgG by copper free click chemistry was achieved in three steps. In the first step, NDs were prepared from scaffold proteins incorporating a single point mutation to cysteine (D73C). Solvent exchange of ND samples to HEPES 20mM NaCl 150mM pH 7.2 was performed by gel filtration using Zeba spin columns (40 kDa MWCO) followed by filtration using a 0.22 μm PES syringe filter. Freshly prepared TCEP stock solution was diluted to a final ×10 fold of MSP and the solution was stirred for 30 minutes at room temperature to achieve complete reduction of cysteines. A bifunctional linker, Mal-PEG(24)-DBCO dissolved in DMSO was added to a final x50 fold over MSP and the mixture was allowed to react for 1.5 hours at room temperature, stirred in the dark. Excess reactants were removed by gel filtration using zeba columns and the sample was fractionated on a size exclusion column to remove residual reactants that could interfere with further steps. Collected sample was concentrated using an Amicon ultrafiltration device (50 kDa MWCO) to a final ND concentration of ~30 μM. The degree of labeling by the linker was evaluated by the molar extinction coefficient of the labeled product to be ~1.2:1 linker:ND (Figure S3A and S3B). In the next step, a IgG polyclonal anti-GABA-A antibody was modified to contain an azide on the Fc using a dedicated kit (ClickSite). Finally, ND-DBCO and Ab-azide were mixed to yield a 2:1 Ab:ND solution (20 μM MSP) and incubated over night at 35 °C in the dark. Sample composition was evaluated by size exclusion chromatography (Fig. S3C).

### Image acquisition

A 100X oil-immersion objective (UAPON100XOTIRF) an EMCCD camera (iXon Ultra 897, Andor) coupled to an inverted microscope (IX83, Olympus), fitted with both epifluorescence and TIRF illumination optics, were used for all imaging experiments. An LED light source (SPECTRA X, Lumencor) was used for epifluorescence illumination and a laser (iChrome MLE, Toptica Photonics) for TIRF illumination.

### Time-resolved fluorescence measurements

Fluorescence lifetime measurements of labeled NDs constructs were performed in a time-correlated time-tagged recording mode using a static confocal excitation and detection spot in the solution. A 470 nm diode laser (LDH-P-C-470B, PicoQuant), connected to a picosecond laser driver (PDL 828 Sepia II, PicoQuant), was coupled to an inverted confocal microscope (SP8, Leica), using a 63X water-immersion objective (HC PL APO 63x/1.20 W, Leica). The emitted light was spectrally filtered at 520±15 nm and 700±30 nm for TF and Cy5, respectively, and focused onto photodetectors (HyD, Leica). The laser driver and photodetectors were connected to a dedicated time-correlated single photon counting system (PicoHarp 300, PicoQuant) in order to obtain lifetime curves. ND samples were diluted to a final 30nM in PBS and transferred to chambered coverslips for measurement. Lifetime measurements in the presence of DPA were performed on the same well by dilution x200 of a stock solution of DPA to a final concentration of 2uM.

### Photobleaching experiments

#### Coverslip preparation

Measurements were performed on NDs adsorbed to chambered coverslips (μ-slide 8 well, Ibidi). Chambers were cleaned by overnight incubation with a 2% Hellmanex III cleaning solution at RT, followed by extensive wash with DDW and 15 minutes sonication in water to remove residual detergent. Finally, residual fluorescent contaminants were removed by a glow discharge system (Emtech K100x, Quorum Technologies) for a total of 3 min.

#### Photobleaching measurements

ND samples were diluted ×10^5^ in PBS to ~30 pM, and 200 μl of the diluted solution was added to each well. Photobleaching measurements of all samples were performed on the same day. Samples were incubated for 10 minutes in the dark at room temperature and the wells were washed six times with PBS. Imaging was performed in a TIRF configuration, using a 488 nm laser line (iChrome MLE, Toptica Photonics) with a total illumination power of 1 mW after the objective. Fluorescence emission of TF and Cy5 were spectrally filtered using a 520±15 nm and a 700±30 nm bandpass filters (Chroma), respectively. Image recording was done with 100 ms exposure time and 300x electron multiplier gain. In order to obtain complete emission curves, not degraded by photobleaching that occurs while focusing the image, we took the following steps: (1) the laser beam is focused on a selected ROI, (2) beam shutter is closed, (3) stage moves ~100 μm in either direction, (4) data acquisition is initiated and finally (5) the beam shutter is opened to collect emission data. While the system was relatively stable, switching between ROIs often destroyed the focus and therefore in a typical experiment it took 2-3 attempts before a stable focus was maintained between ROIs.

#### Data analysis of photobleaching measurements

we used PIFit, a dedicated software for data analysis of photobleaching measurements using a TIRF setup. In the analysis we used a 15% intensity threshold for spot detection, a 2σ limit for proximity between spots and a threshold of 2.5 step/noise. Analysis parameters were identical for all datasets. These parameters were calculated using the PIFit software package[21].

### HEK293 and primary cortical neuron culture

Human embryonic kidney (HEK293) cells were cultured in Dulbecco’s Modified Eagle Medium (DMEM) containing 10% fetal calf serum, 2 mM glutamine, 100 units/ml penicillin G and 100 μg/ml streptomycin, and grown in 5% CO_2_ at 37 °C under 90-95% humidity. For electrophysiology experiments the cells were seeded on glass coverslips (25-mm diameter) that were placed in 6-well plates and pre-coated with poly-D-lysine. Electrophysiology experiments were performed 24-48 hours after seeding. Cells were plated at a 1:5 or 1:10 dilution onto poly-D-lysine coated 6-well 3–5 days before an experiment. Experiments were performed with 70% confluent cells. All animal experiments were approved by the local ethics committee for animal research (ethical approval number 63-09-2018). Primary cortical neurons were cultured from P1 and P2 newborn Sprague Dawley rats. The tissue was digested by 100 units of papain (Sigma-Aldrich) in Ca^2+^/Mg^2+^ free Hank’s balanced salt solution (HBSS) (Biological Industries, Israel) in 37 °C incubator. The tissue was then mechanically dissociated, and the cells were plated on 50 μg/ml of poly-D-lysine-coated cover glasses in Neurobasal medium (Biological Industries, Israel) supplemented with 2% B-27 (Invitrogen), GlutaMAX-I 2mM (Invitrogen), antibiotics Penicillin (100 unit/ml) / streptomycin (100ug/ml) (Beit Haemek 030311B), glucose and 5% normal horse serum (Biological Industries, Israel). On the following day and twice a week thereafter, medium was exchanged with growth medium, once a week the medium was applied with Ara-C to prevent glial proliferation. Cell were incubated at 37 °C in humidified air containing 5% CO_2_.

### Colocalization, staining & transfection

The following antibodies were used: GABA-A receptor γ 2 (Synaptic System), as Secondary antibodies, Alexa Fluor 647, Alexa Fluor 546 and Alexa Fluor 488 were used (Invitrogen). Cortical neuron cultures were prepared and grown on glass coverslips (25 mm diameter) placed in were placed in 6-well plates and pre-coated with poly-D-lysine. Cells were transfected after 9 days in vitro with pmCherryC2-Gephyrin P1, a gift from Shiva Tyagarajan (Addgene plasmid # 68820; http://n2t.net/addgene:68820; RRID: Addgene_68820). Transfection was performed using Lipofectamine 3000 (Invitrogen) according to the manufacturer’s instructions (12 μl of Lipofectamine 3000 and 2.8 μg were used per well). Expression was confirmed after 24 hours by visual inspection and fluorescence microscopy. ND_TF,Cy5_ conjugated primary antibody GABA-A receptor γ 2 (Synaptic System) was diluted in fresh cell culture medium. The diluted antibodies were applied directly to the well containing the live cells 24h after fluorescents protein expression, and were incubated for 10 minutes at 37 °C in a 5% CO_2_ incubator. Subsequently, the cells were washed twice with neuronal external solution.

### Patch-clamp experiments

#### Electrophysiological recording

Whole-cell voltage-clamp recordings in HEK cells were performed using a computer-controlled amplifier (EPC 10 USB, HEKA Elektronik). Recorded signals were filtered at 10 kHz using a Bessel filter, and digitized at 10 kHz to 100 kHz, depending on the timescale of the voltage waveform.

The external solution for HEK cells contained (in mM): 140 NaCl, 2.8 KCl, 2 CaCl_2_, 1 MgCl_2_, 10 HEPES and 10 glucose, adjusted with NaOH to pH 7.4 (310 mOsm). The pipette solution contained (in mM): 125 K-Gluconate 125, 0.6 MgCl_2_, 0.1 CaCl_2_, 1 EGTA, 10 HEPES, 4 Mg-ATP, 0.4 Na_2_GTP, adjusted with KOH to pH7.4 (295 mOsm). The electrode resistance was 5–10 MΩ when filled with the pipette solution.

For neural cells all imaging and electrophysiology experiments were conducted with a neuronal external solution containing (in mM): 116.9 NaCl, 2.7 KCl, 1.8 CaCl2, 0.5 MgCl2, 20 HEPES, 5.5 glucose, 0.36 NaH2PO4, 200 Ascorbic Acid and Pen (100 unit / ml)/Strep (100μg/ml) at pH 7.3, and adjusted to 320 mOsm with sucrose. The pipette solution contained (in mM): 110 potassium gluconate, 10 KCl, 10 glucose, 8 Na creatine-phospate, 5 EGTA, 10 HEPES, 4 Mg-ATP, 0.4 Na-GTP (pH 7.3); adjusted to 290 mOsm with sucrose.

#### Optical recording

Imaging was performed using an LED excitation (SPECTRA X, Lumencor) passing through a 470/24 nm bandpass filter, and an EMCCD camera (iXon Ultra 897, Andor), coupled to a motorized inverted microscope (IX83, Olympus). All imaging was performed using a 100X oil-immersion objective (UAPON100XOTIRF). Fluorescence emission was filtered with a bandpass filter centered at 590 nm for mCherry (ET590/33m, Chroma), and centered at 700 nm for Cy5 (ET700/75m, Chroma). A DAQ device (USB-6211, National Instruments), controlled by a home-written software (LabVIEW, National Instruments), was used for synchronizing the camera and the patch clamp amplifier

#### Data analysis

Analysis of fluorescence imaging of clamped cells was performed in MATLAB (MathWorks). For sparse staining, a 2D particle tracking algorithm was used in order to correct for small translations. Fluorescence intensity for each frame was calculated by averaging the 10 brightest pixels of each spot for sparse staining, and averaging the fluorescence from a whole cell for ensemble measurements. For the latter, background-subtraction of each frame was performed by subtracting the mean value of the 10% darkest pixels. The average optical voltage response for each intensity trace was calculated by averaging the intensity value corresponding to every voltage level. In order to achieve a stabilized value of the response, the averaging was performed on the frame containing the last 100 ms of each voltage step.

