## Supplementary material for "Point-localized and site-specific membrane potential optical recording by engineered single fluorescent nanodiscs": SI - Point-localized and site-specific membrane potential optical recording by engineered single fluorescent nanodiscs

### Supplementary information figures:

Figure S1:

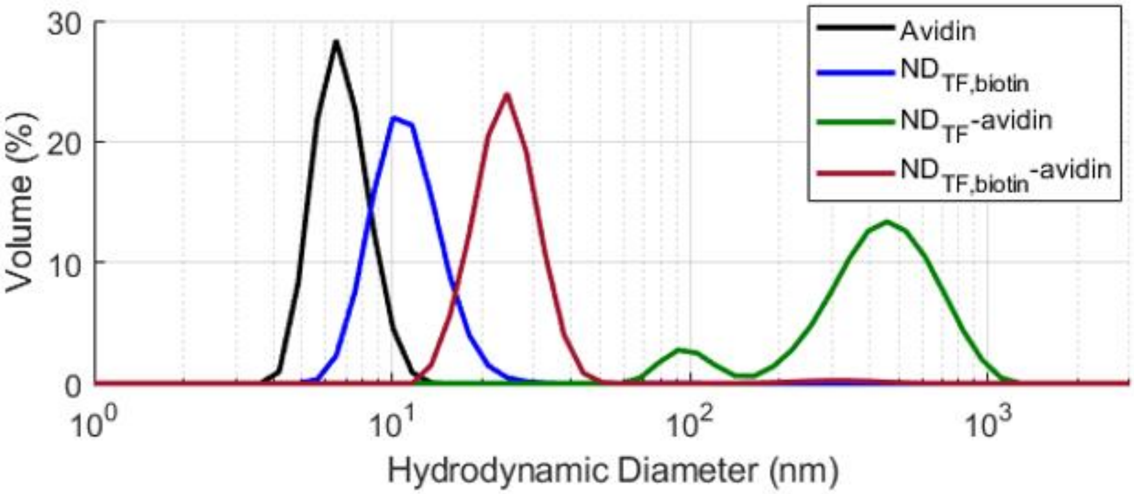


**Figure S1.** The effect of avidin binding to NDs demonstrated by DLS measurements: The hydrodynamic diameter of free avidin in solution is 7.4±1.3 nm and for free ND in solution is 13.1±2.5 nm. Binding of avidin to ND-biotin results in a single complex with a hydrodynamic diameter of 27.2±4.6 nm after 20 min, exactly matching a construct of one ND with avidin on both facets (13 + 2×7.5, most likely more than one avidin on each facet). In contrast, ND without biotin in its lipid composition incubated with avidin results in two populations of large sizes. This is the result of nonspecific binding between the positively charged avidin and the negatively charged ND. This further supports the specificity of ND-biotin binding to avidin.

Figure S2:

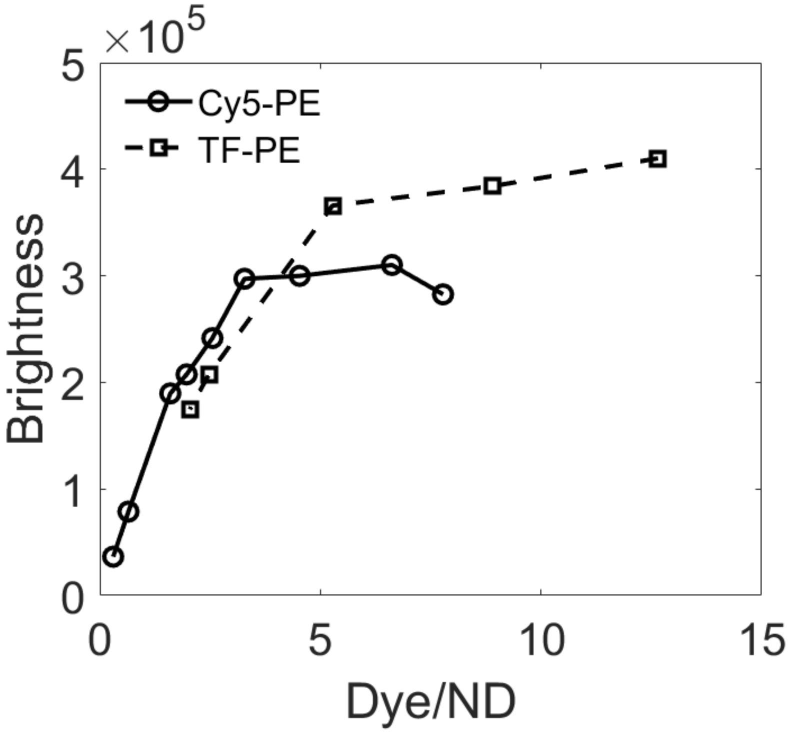


**Fig. S2.** Brightness of NDs incorporating TF-PE and Cy5-PE. The brightness is the product of the QY, the average number of dyes per ND and the apparent extinction coefficient.

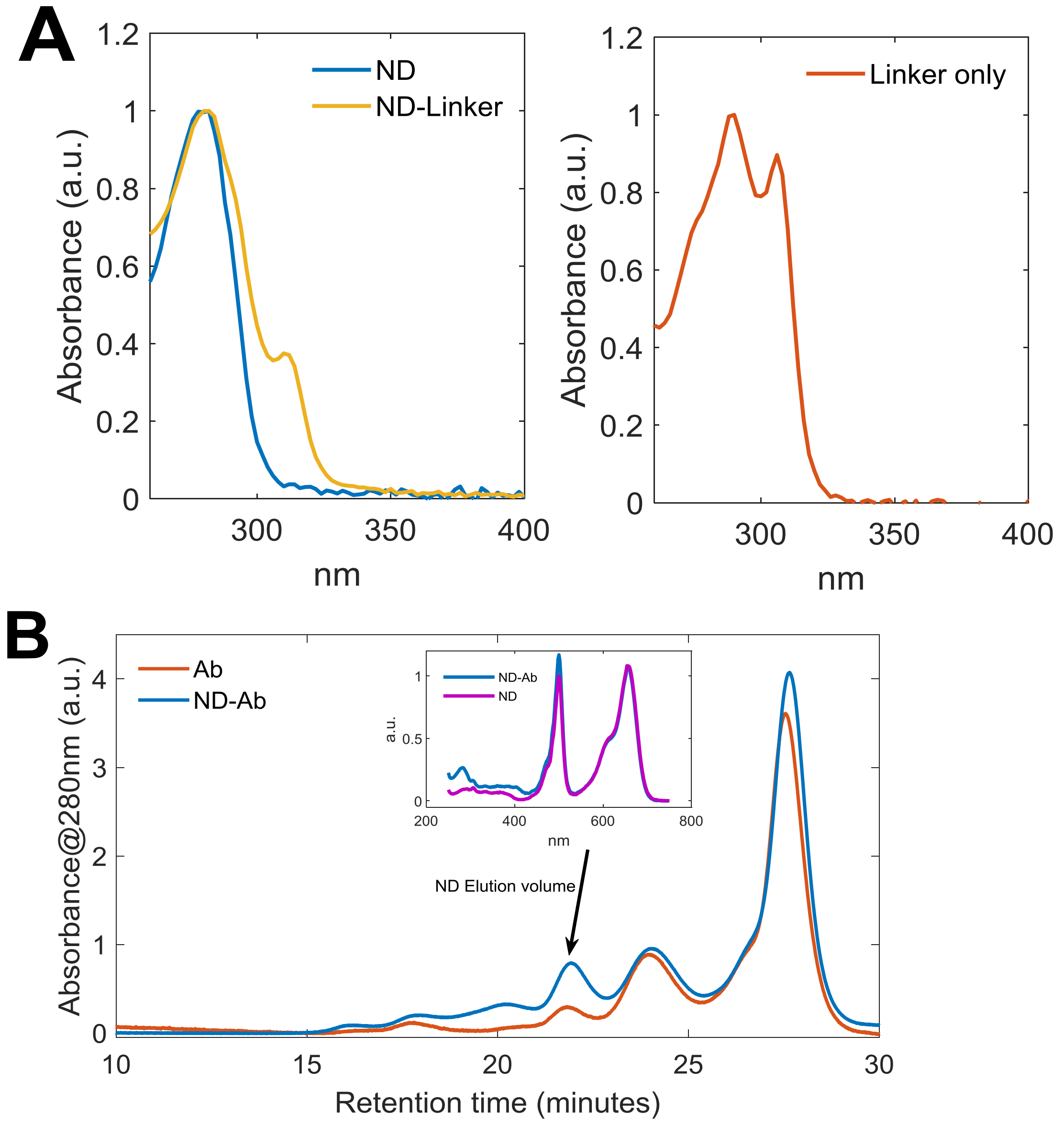


**Fig. S3.** Validation of Ab crosslinking to ND. (A) The degree of labeling for the Mal-PEG$_{24}$-DBCO linker to cysteine groups on MSP was evaluated by absorption spectra analysis. The contribution of the linker to the absorbance at 280nm for the ND-linker conjugate was determined from the spectra of the free linker to be 80% of the linker absorbance at 309nm. The relative concentrations of MSPs and linker [MSP]=24uM and [linker]=29uM leading to a labeling ratio of ~1:1.2 ND:linker. (B) Ab conjugation to labeled NDs is confirmed by an increase in the UV spectra of the labeled NDs. the conjugated ND-Ab fraction was collected and the purified product was further concentrated using a 50kD ultrafiltration device and immediately used for cell labeling.

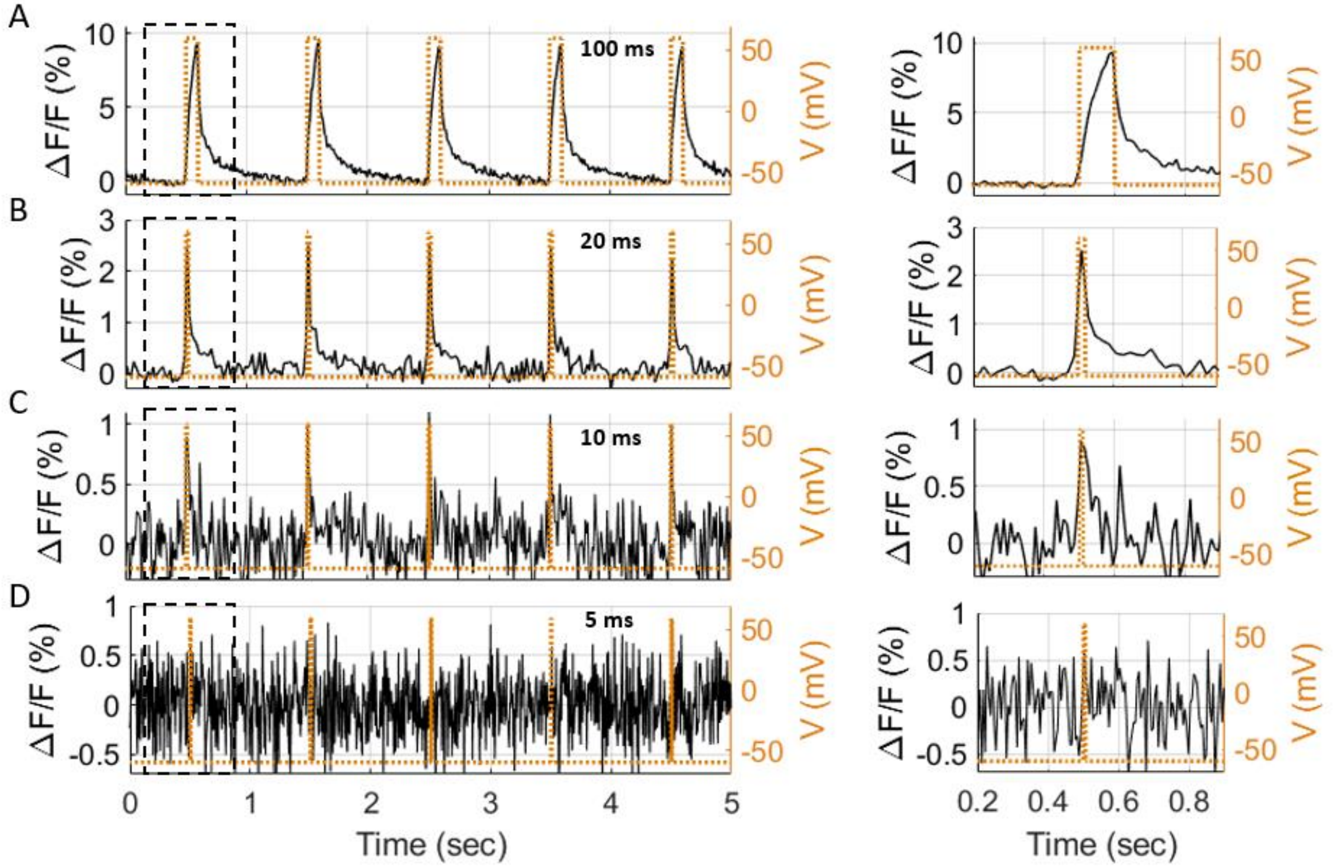


**Fig. S4.** Response kinetics of $ND_{TF,Cy5}$ – DPA to changes of membrane potential in a HEK293 cells. Cells were stained with $ND_{TF,Cy5}$, and voltage-clamped in the presence of 2 μM DPA. Cells were voltage-clamped to a waveform consisting of 120 mV pulses with varying durations, and a -60 mV resting potential. Left figures show the fluorescence response of Cy5 of a representative cell with pulse durations of **(A)** 100 ms, **(B)** 20 ms, **(C)** 10 ms, and **(D)** 5 ms. Right figures show zoom-in on the duration indicated by the dashed square.

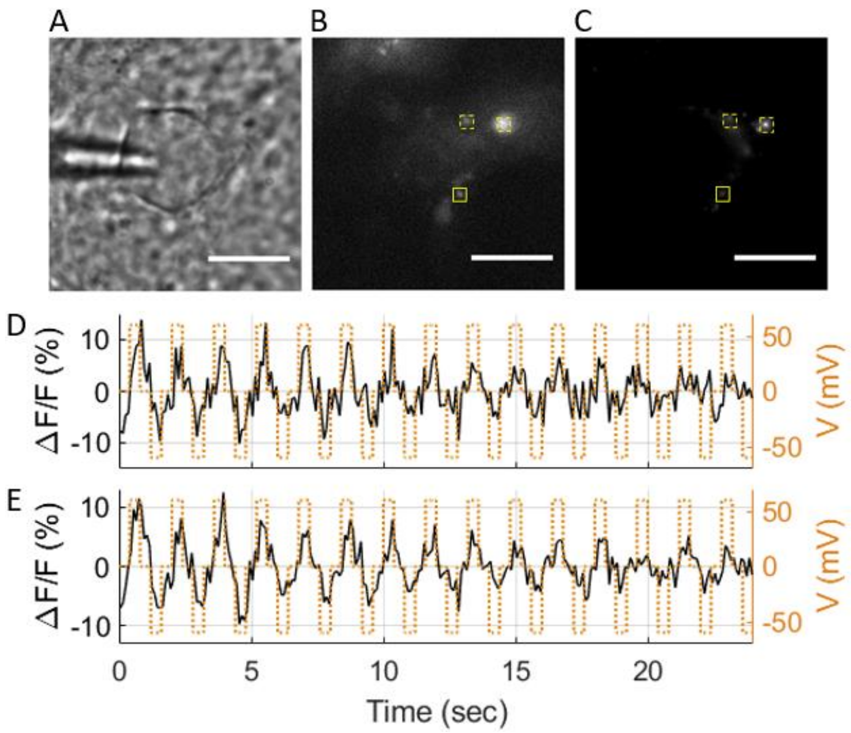


**Fig. S5.** Single particle membrane potential optical recordings from $ND_{TF,Cy5}$-Ab-avidin complex targeting $GABA_AR$, and DPA. Cultured cortical neurons were transfected with mCherry-gephyrin and stained with $ND_{TF,Cy5}$ conjugated to Ab-GABA and avidin. Cells were voltage-clamped in the presence of 2 µM DPA. **(A)** DIC image of a patched cortical neuron, expressing mCherry-gephyrin and labeled with $ND_{TF,Cy5}$-Ab-avidin. **(B)** and **(C)** show the emission of mCherry and Cy5, respectively. Scale-bar: 10 µm. **(D)** Shows the changes in Cy5 fluorescence emission as a function of membrane potential from the $ND_{TF,Cy5}$ inside the solid yellow square. **(E)** Shows the change in Cy5 fluorescence emission as a function of membrane potential from the $ND_{TF,Cy5}$ averaged from the three NDs inside indicated by yellow squares. The average responses of the single and averaged response trajectories were 8.9±7% and 9.7±5.5% per 120 mV, respectively,

.
